# Overexpression of α-SNAP_Rhg1_ can improve *rhg1-a* mediated soybean resistance to soybean cyst nematode

**DOI:** 10.1101/2024.05.18.594834

**Authors:** Deepak Haarith, Soumita Das, Emma Nelson, Ryan Zaptocony, Andrew Bent

**Author notes:** Department of Plant Pathology, University of Nebraska, Lincoln, NE 68583, USA. School of Integrative Plant Science, Cornell University, Ithaca, NY 14853, USA. PPD/ThermoFisher Scientific, Middleton, WI 53562.

## Abstract

The *rhg1-a* and *rhg1-b* haplotypes of the soybean *Rhg1* locus are economically effective tools for the control of soybean cyst nematode (SCN; *Heterodera glycines*), but ongoing SCN evolution requires improved sources of resistance. Both *Rhg1* haplotypes carry multiple tandem repeat copies of a four-gene block encoding four disparate proteins; resistance efficacy scales with copy number. The haplotypes encode different variants of an unusual α-SNAP protein whose abundance increases in the nematode-reprogrammed plant cells that form the syncytium (nematode feeding site), which subsequently collapses. Simultaneous presence of the two α-SNAP_Rhg1_ protein types, or substantially elevated expression of either type, was hypothesized to improve SCN resistance but has not been achievable by conventional plant breeding methods. We accomplished both, by transgenic additions to an *rhg1-a Rhg4* soybean line. Existing resistance to HG type 0 and HG type 2.5.7 SCN was strengthened, but resistance was not improved against an HG type 1.3.6.7 SCN population that was already highly virulent on the parent line. No significant yield penalties from constitutive overexpression of transgenic α-SNAP genes were observed in greenhouse or initial field studies. Cisgenic combination or elevated expression of the different α-SNAP genes may extend the efficacy and/or durability of *Rhg1*-mediated resistance. We also observed differential protein abundances for some α-SNAP_Rhg1_ isoforms after inoculation with different HG type populations of SCN. These approaches provide opportunities to understand the different mechanisms of the two *Rhg1* haplotypes and to possibly fine-tune them for better management of SCN.

## (ii) Introduction

Vesicular transport is a salient feature of all eukaryotic cells and is necessary for cellular homeostasis. Two proteins, α-SNAP (α-Synaptosomal NSF Associated Protein) and NSF (N-ethylmelamide Sensitive Factor), are crucial for ongoing vesicular transport because they disassemble the SNARE protein bundles that form from v-SNAREs and t-SNAREs during vesicle docking to target membranes {Huang, 2019 #96}. As an allopolyploid, the soybean genome encodes at least five α-SNAP proteins, including one encoded at the *Rhg1* locus (*resistance to <u>H</u> eterodera <u>g</u>lycines*) on chromosome 18 (Cook et al., 2012; Lakhssassi et al., 2017). *Rhg1* is a strong-effect quantitative trait locus that contributes to resistance against soybean cyst nematode (SCN, *Heterodera glycines*), the most economically damaging disease of soybean (Bent, 2022; Bradley et al., 2021; Concibido et al., 1997). SCN resistance-conferring isoforms of the α-SNAP_Rhg1_ proteins encoded by *Glyma*.*18G022500* (also known as *GmSNAP18*) are unusual because they carry divergent amino acids at the six C-terminal amino acids of α-SNAP, a key α-SNAP/NSF interaction site that is otherwise highly conserved across Eukaryota (Cook et al., 2014). Soybean plants that carry SCN resistance-conferring α-SNAP_Rhg1_HC or α-SNAP_Rhg1_LC proteins are not viable if they do not also carry a unique NSF isoform, NSF_RAN07_, which contains amino acid changes that allow improved binding between NSF and the α-SNAP_Rhg1_HC or α-SNAP_Rhg1_LC proteins (Bayless et al., 2018).

*Rhg1* is a complex locus that exhibits tandem repeat copy number variation of a 31 to 36 kb genome segment encoding four distinct genes, three of which have been shown to contribute to SCN resistance (Bayless et al., 2019; Cook et al., 2012; Dong & Hudson, 2022; Liu et al., 2017). Soybean germplasm was originally dominated by a single-copy *Rhg1* haplotype now known as *rhg1-c* (Lee et al., 2015). SCN resistance breeding has expanded the presence of two additional major haplotypes, lower-copy-number *rhg1-a* (exemplified by the three-copy haplotype from the Peking soybean accession that encodes α-SNAP_Rhg1_LC), and higher-copy number *rhg1-b* (exemplified by the ten-copy haplotype from the PI 88788 soybean accession that encodes α-SNAP_Rhg1_HC) (Bent, 2022; Mitchum, 2016). The *rhg1-b* locus has worked well on its own to confer SCN resistance and over 95% of SCN-resistant soybean varieties now marketed in the U.S. carry *rhg1-b* (e.g.,Tylka et al., 2021). However, recurrent planting of *rhg1-b* soybeans is, over multiple years, leading to the emergence of local SCN populations that partially but increasingly overcome the resistance mediated by *rhg1-b* (Kleczewski et al., 2023; McCarville et al., 2017; Niblack et al., 2008). The *rhg1-a* haplotype is less effective on its own, but highly effective when coupled with an enzyme-altering allele of the Serine Hydroxy-Methyl Transferase *Rhg4* gene Glyma.08g108900 on chromosome 8 (*GmSHMT08*, encoding SHMT_*Rhg4*_), and/or loss-of-function alleles of two other α-SNAP-encoding loci, *Rhg2* (*GmSNAP11*) on chromosome 11 or *GmSNAP02* on chromosome 2 (Basnet et al., 2022; Liu et al., 2012; Meksem et al., 2001; Patil et al., 2019; Shaibu et al., 2022; Usovsky et al., 2023). There is also evidence for physical interaction of the α-SNAP_Rhg1_LC and SHMT_*Rhg4*_ proteins (Lakhssassi, Piya, Knizia, et al., 2020).

At present we have limited understanding of how the *Rhg1*-encoded proteins AAT_Rhg1_ and WI12_Rhg1_ contribute to SCN resistance, but more has been learned about α-SNAP_Rhg1_ (Bent, 2022; Dong & Hudson, 2022; Han et al., 2023). Apparently, disease resistance arises through alteration of α-SNAP/NSF interaction and vesicular trafficking caused by the aberrant *Rhg1* α-SNAPs (Bayless et al., 2016). Relative to the surrounding root cortical cells, the level of α-SNAP_Rhg1_HC or α-SNAP_Rhg1_LC protein is elevated ∼15-fold in syncytial plant root cells that are reprogrammed by nematode effectors to act as nematode feeding sites. Syncytia constitute the obligatory biotrophic interface where SCN must feed for three to four weeks but in *rhg1*-mediated resistance, syncytial collapse is observed within a few days of syncytium initiation (Mitchum, 2016).

The continual emergence of SCN populations that can overcome *rhg1-b* or *rhg1-a*/*rhg2*/*Rhg4*-mediated resistance threatens soybean production and motivates exploration of improved approaches for SCN resistance. Different populations of SCN evolve virulence against different sources of resistance and are classified as distinct HG types (Niblack et al., 2006). While HG 0 SCN populations are considered avirulent on either *rhg1-a* (Peking type) or *rhg1-b* (PI 88788 type), HG 1.3.6 populations are at least partially virulent on *rhg1-a Rhg4* plants, and HG 2.5.7 populations are at least partially virulent on *rhg1-b* plants (Niblack et al., 2002). HG 2.5.7 populations are the more immediate issue in the U.S. areas with the highest soybean production, due to heavy use of *rhg1-b* (Kleczewski et al., 2023; McCarville et al., 2017). Combining *rhg1-b* with smaller-effect SCN resistance QTL, none of which give strong resistance on their own, has been shown to restore valuable levels of resistance against HG 2.5.7 and other SCN (Brzostowski & Diers, 2017). Most SCN-resistant germplasm accessions studied to date carry identical or highly related variants of *rhg1-a* or *rhg1-b* (Cook et al., 2014; Lee et al., 2015). Published searches for functionally novel *Rhg1* haplotypes have to date identified only one qualitatively novel haplotype, the apparent G. soja progenitor of *rhg1-a* and *rhg1-b*, whose utility is not certain (Grunwald et al., 2022). One alternative is to use cisgenic transgene technologies to generate lines that express amounts, types and/or combinations of soybean *Rhg1* proteins that have not been obtainable through conventional plant breeding.

In the present study, we generated and characterized soybean lines that overexpress either the *rhg1-a* type α-SNAP (α-SNAP_Rhg1_LC) or *rhg1-b* type α-SNAP (α-SNAP_Rhg1_HC) in the elite public soybean varieties IL3849N (*rhg1-a* genetic background) and IL3025N (*rhg1-b* genetic background). The resistance of the resulting lines against different SCN HG types was assessed. Changes in transcript and/or protein abundance in response to infections by different SCN HG types were monitored for α-SNAP_Rhg1_HC, α-SNAP_Rhg1_LC, NSF and a limited set of defense-associated indicator genes. A third objective was to evaluate the effect of constitutively overexpressing potentially toxic rhg1 genes on quantitative and qualitative aspects of seed production. Quantitative enhancements of SCN resistance were observed, suggesting that cisgenic modification of rhg1 α-SNAP expression and/or type merits further investigation as a means of improving SCN resistance in soybean.

## (iii) Results

### Soybean lines with overexpression of α-SNAP_Rhg1_LC or α-SNAP_Rhg1_HC

We obtained transgenic lines that overexpress either the *rhg1-a* α-SNAP (α-SNAP_Rhg1_LC) or the *rhg1-b* α-SNAP (α-SNAP_Rhg1_HC) under the control of a soybean ubiquitin promoter in high-yielding public sector soybean varieties. We used two different host soybean lines for transformation: IL3849N was originally bred to carry *rhg1-a* as well as *Rhg4*, and IL3025N was originally bred to carry *rhg1-b*. A table summarizing these lines and a flow chart for the present study are provided in Figure S1. Several independent transformants (events) were obtained for α-SNAP_Rhg1_LC in the IL3849N (low-copy) background, but no transformants were obtained for α-SNAP_Rhg1_LC in the IL3025N (high-copy) background. For α-SNAP_Rhg1_HC we obtained only one overexpressing transformant per line in both soybean backgrounds. The resulting lines are not cisgenic lines because they utilize an *Agrobacterium nos* terminator downstream of the *GmSNAP18* gene and a heterologous 35S:aadA spectinomycin resistance selectable marker, but they model potential cisgenic or intragenic overexpression of soybean α-SNAP_Rhg1_ proteins in soybean (Schouten & Jacobsen, 2008).

### Overexpressing either α-SNAP_Rhg1_HC or α-SNAP_Rhg1_LC in IL3849N (*rhg1-a*/*Rhg4*) enhanced resistance against HG 0 and HG 2.5.7 SCN populations

We determined the relative SCN resistance of the resulting overexpression IL3849N lines by quantifying the FI (Female Index) of SCN HG 0 IL, HG 2.5.7 IL and HG 1.3.6.7 MN 30 days after inoculation. The IL3849N background is already resistant to both HG 0 IL and HG 2.5.7 IL (FI < 10). Readers are reminded that HG 0 SCN have low virulence on *rhg1-a* (Peking type) and *rhg1-b* (PI 88788 type) plants, HG 1.3.6.7 populations are partially virulent on *rhg1-a* plants (IL3849N), and HG 2.5.7 populations are partially virulent on *rhg1-b* plants (IL3025N). Overexpressing α-SNAP_Rhg1_LC or α-SNAP_Rhg1_HC in IL3849N background (subsequently referred to as IL3849N+HC and IL3849N+LC, respectively) significantly decreased the FI of the HG 0 IL and HG 2.5.7 IL SCN populations (Figure 1a and 1b). Although for both LC and HC overexpression in the IL3849N background, the reduction of FI for HG 2.5.7 IL was statistically significantly in at least one biological replicate, only overexpression of α-SNAP_Rhg1_HC in the LC background exhibited a significant enhancement of resistance across both experiments (Figure 1b). However, no significant reduction of FI was observed for HG 0 IL or for HG 2.5.7 IL in IL3025N lines overexpressing HC α-SNAP_Rhg1_ (IL3025N+HC) (Figure S2a, b).

**Figure 1.**
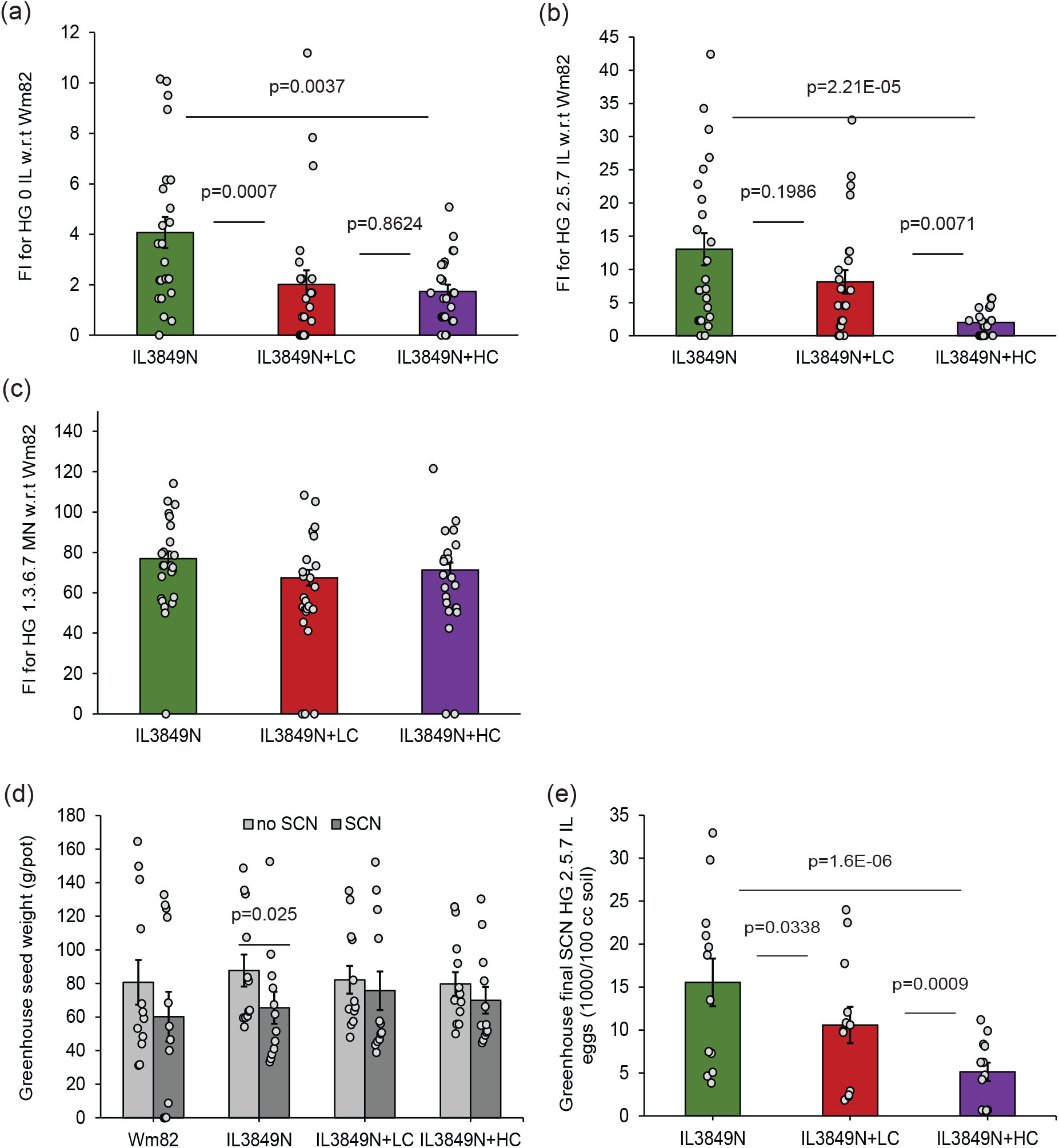
SCN reproduction, seed yield and end-of-season SCN egg counts for IL3849N transgenic lines. Female indices (FI, SCN cyst formation relative to the SCN-susceptible control line Wm82) for parent IL3849N (*rhg1-a*) line compared to IL3849N+LC and IL3849N+HC, for (a) SCN HG 0 IL (b) SCN HG 2.5.7 IL and (c) SCN HG 1.3.6.7 MN. All FI experiments were done twice (Set 1 and Set 2 seeds) with 12 replicates each in a randomized design. (d) Greenhouse seed yield for Wm82, IL3849N, IL3829N+LC and IL3849N+HC. The greenhouse experiment was repeated three times (using Set 1, Set 1, and Set 2 seeds), with four randomized complete block replicates and surrounding IL3849N border in each experiment. (e) Comparison of end-of-season SCN egg density (eggs/100 cc soil) for IL3849N, IL3849N+LC and IL3849N+HC. Error bars show standard error of the mean; p-values are for ANOVA Tukey HSD.

### Overexpressing either α-SNAP_Rhg1_HC or α-SNAP_Rhg1_LC failed to significantly enhance resistance against an HG 1.3.6.7 SCN

When the above IL3849N α-SNAP_Rhg1_ overexpression lines were challenged with an HG 1.3.6.7 MN population that is already virulent on the IL3849N background (FI > 55), overexpression of either the HC or LC α-SNAP_Rhg1_ (IL3849N+HC and IL3849N+LC respectively) failed to significantly reduce the female index (Figure 1c). However, a slight trend of reduction in FI was observed for the cis-genics compared to the parent IL3849N LC background (Figure 1c). We also obtained a single transgenic event for IL3025N (*rhg1-b*) overexpressing α-SNAP_Rhg1_HC (termed IL3025N+HC), and no significant reduction of FI was observed when HG 1.3.6.7 MN SCN were inoculated that line (Figure S2c).

### Overexpressing α-SNAP_Rhg1_HC in IL3025N (*rhg1-b*) did not significantly enhance resistance

For IL3025N+HC, similar to the results with HG 1.3.6.7 MN SCN (see above), no improvement of resistance against HG 0 IL or HG2.5.7 IL SCN populations was detected (Figure S2).

### Overexpressing α-SNAP_Rhg1_LC or α-SNAP_Rhg1_HC in IL3849N reduces reproduction of SCN HG 2.5.7 IL without detectably compromising yield

Three randomized complete block design experiments were conducted in a greenhouse (four blocks per experiment) to further test SCN reproduction and also monitor the seed yield of the transgenic lines. In the first trial (experiment 1, Winter 2020), three out of four SCN-susceptible control Williams 82 (Wm82) plants inoculated with SCN died at around 40 dpi. No plant deaths were observed for the second (experiment 2, Summer 2021) and the third (experiment 3, Winter 2021) trials. The plants grew larger and produced higher seed yield across all treatments during the summer 2021 experiment compared to the Winter 2020 and Winter 2021 experiments despite supplemental winter heating and lighting in the greenhouse. All greenhouse trials used HG 2.5.7 IL SCN. Transgenic disease resistance implementations often carry a yield penalty (Collinge et al., 2010). However, we observed no statistically significant differences in the seed yield (total seed weight corrected to 13% moisture) between the transgenic lines and the parental line in this set of greenhouse experiments (light grey bars, Figure 1d). The spread of data for seed yield in Figure 1d reflects the lower seed biomass obtained in the winter trials of 2020 and 2021 relative to the summer 2021 trial, but the ANOVA blocked by experiments focuses on within-experiment variation between genotypes, which was not significant (Table S1).

When comparing greenhouse seed yield between plants inoculated with SCN HG 2.5.7 IL and non-inoculated plants of the same genotype, only the IL3849N parental line showed a significant drop in seed mass yield across the three experiments (Figure 1d).

IL3849N and the IL3849N transgenic lines did not differ in their seed oil content (Table S1). Oil content of the seed from IL3849N lines also did not change significantly under SCN infection (Table S1). Wm82 had significantly lower oil content compared to IL3849N and the cis-genics, and a Tukey HSD p = 0.049 significance for difference with vs. without SCN infection. No significant differences in seed protein content were observed among IL3849N and the transgenics while Wm82 had significantly less protein overall. SCN inoculation did not change the overall seed protein content in any genotype significantly. There also were no significant differences in seed fiber content between Wm82, IL3849N and the IL3849N transgenics, or with/without SCN (Table S1).

As expected, in all three greenhouse trials, there were far fewer nematode eggs in the soil post-harvest for any of the IL3849N control or transgenic plants compared to Wm82 plants (Figure S2d). IL3849N+HC had substantially lower final nematode egg density compared to the parent IL3849N or IL3849N+LC (Figure 1e). The IL3849N+LC line also had lower end-of-season egg abundance than the parental IL3849N (Tukey HSD p = 0.034). These SCN resistance findings of Figure 1e (greenhouse experiment post-harvest egg count in the soil) were consistent with the findings of the Figure 1b (growth chamber experiments cyst count at 35 days post inoculation).

Permits were subsequently obtained to grow IL3849N and the transgenic IL3849N lines overexpressing α-SNAP_Rhg1_LC or α-SNAP_Rhg1_HC in 2023 in an eastern Nebraska field plot considered to have little or no SCN presence. This single-site, single-year trial used a randomized complete block design of four blocks with 3 m rows (3.3 m until mid-season alley trimming) and 76 cm row spacing. Two adjacent rows of the same genotype were harvested together for each entry. Two different events or lines were used for the IL3849N lines overexpressing α-SNAP_Rhg1_LC or α-SNAP_Rhg1_HC, for a total of five lines per block. All entries were bordered with IL3849N genotypes on all sides. All lines (maturity group 3.8) grew well and had a similar physical appearance, and no statistically significant difference in yields was observed. However, a trend of lower average yield was present in the transgenic lines (Figure S2e). No SCN enumeration was conducted in the fields prior to planting but end-of-season data confirmed that SCN pressure across the utilized research plot was minimal (end-of-season average SCN egg count between 0-80 eggs/100cc soil; no significant differences across the different plant genotypes).

### HG 2.5.7 IL SCN induce plant transcript and protein abundance responses that differ from those induced by HG 0 IL and HG 1.3.6.7 MN

Transcript abundance and corresponding protein abundances were monitored for *GmSNAP18* as well as a small set of additional loci. Many results were as expected but as is described more fully below, the HG 2.5.7 IL SCN population (but not the other SCN populations) often caused surprisingly different regulation of overexpressed α-SNAP_Rhg1_LC (Figure 2, Figure 3, Figure S4).

**Figure 2.**
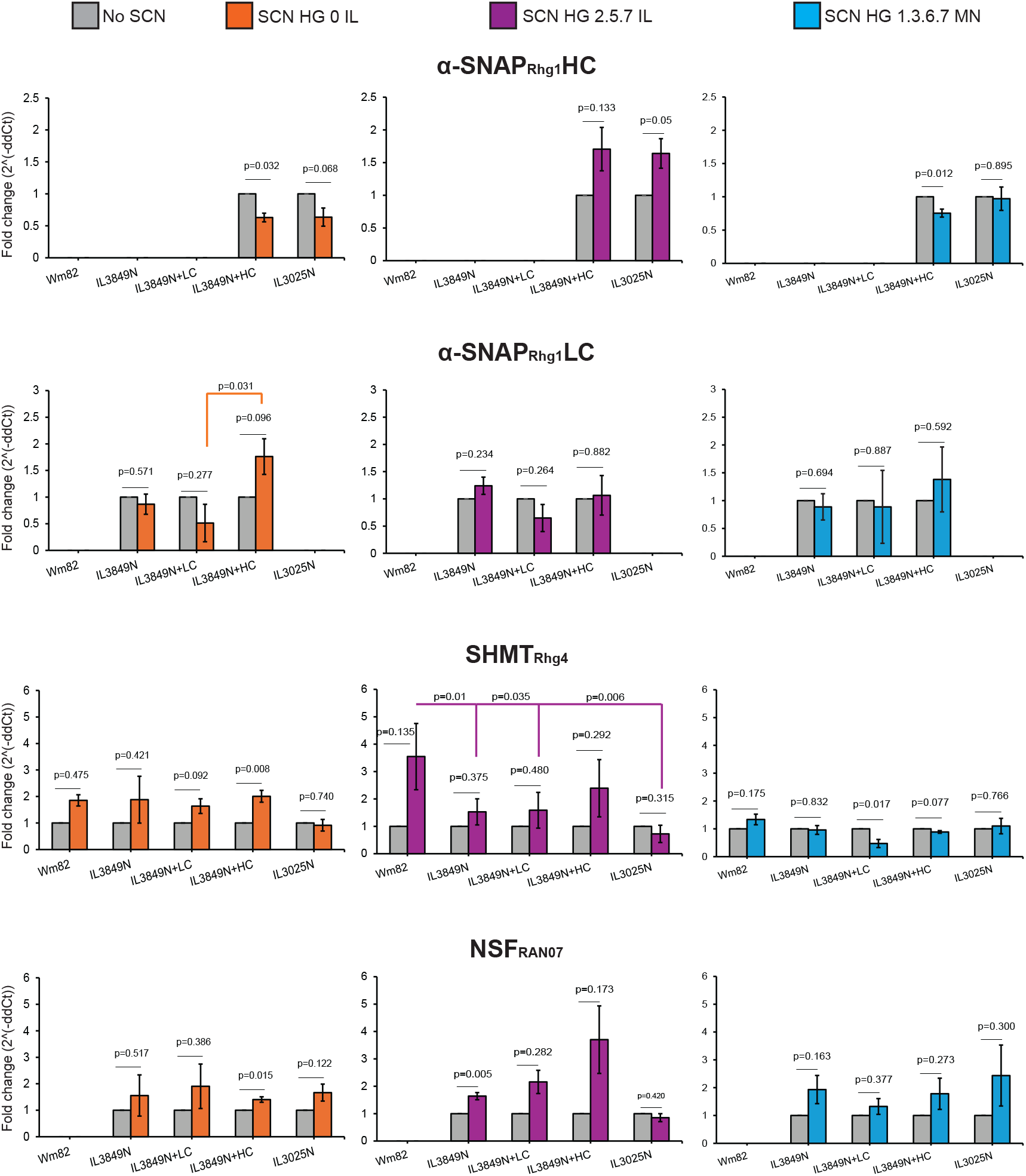
Relative abundance of transcripts encoding α-SNAP_Rhg1_LC, α-SNAP_Rhg1_HC, SHMT_Rhg4_ and NSF_RAN07_, for SCN-infested root zones (colored bars) or similar areas from non-infested samples (grey bars) from the IL3849N transgenic lines and controls. Determined by qPCR with allele-specific primers, using 7 dpi infection-zone root samples for Wm82 (Susceptible), IL3849N (*rhg1-a* parent), IL3849N+LC, IL3849N+HC and IL3025N (*rhg1-b* parent). Transcript fold-change (y-axis) is relative to similar root segments from non-inoculated roots of the same genotype; error bars show standard error of the mean. Statistical p-values above grey horizontal lines compare non-inoculated to inoculated samples of same plant genotype, for each SCN HG type (two-tailed t-tests). qPCR not performed if the target allele is not present in that plant genotype. Statistical p-values above colored bars compare different plant genotypes, and are shown only for p < 0.05 instances indicating that response to the same SCN population differed between the indicated plant genotypes. All experiments done on two separate roots each from Set 1 and Set 2 seedlings (four roots total); each root tested with two qPCR technical replicates. Additional statistical tests shown in Figure S3b.

**Figure 3.**
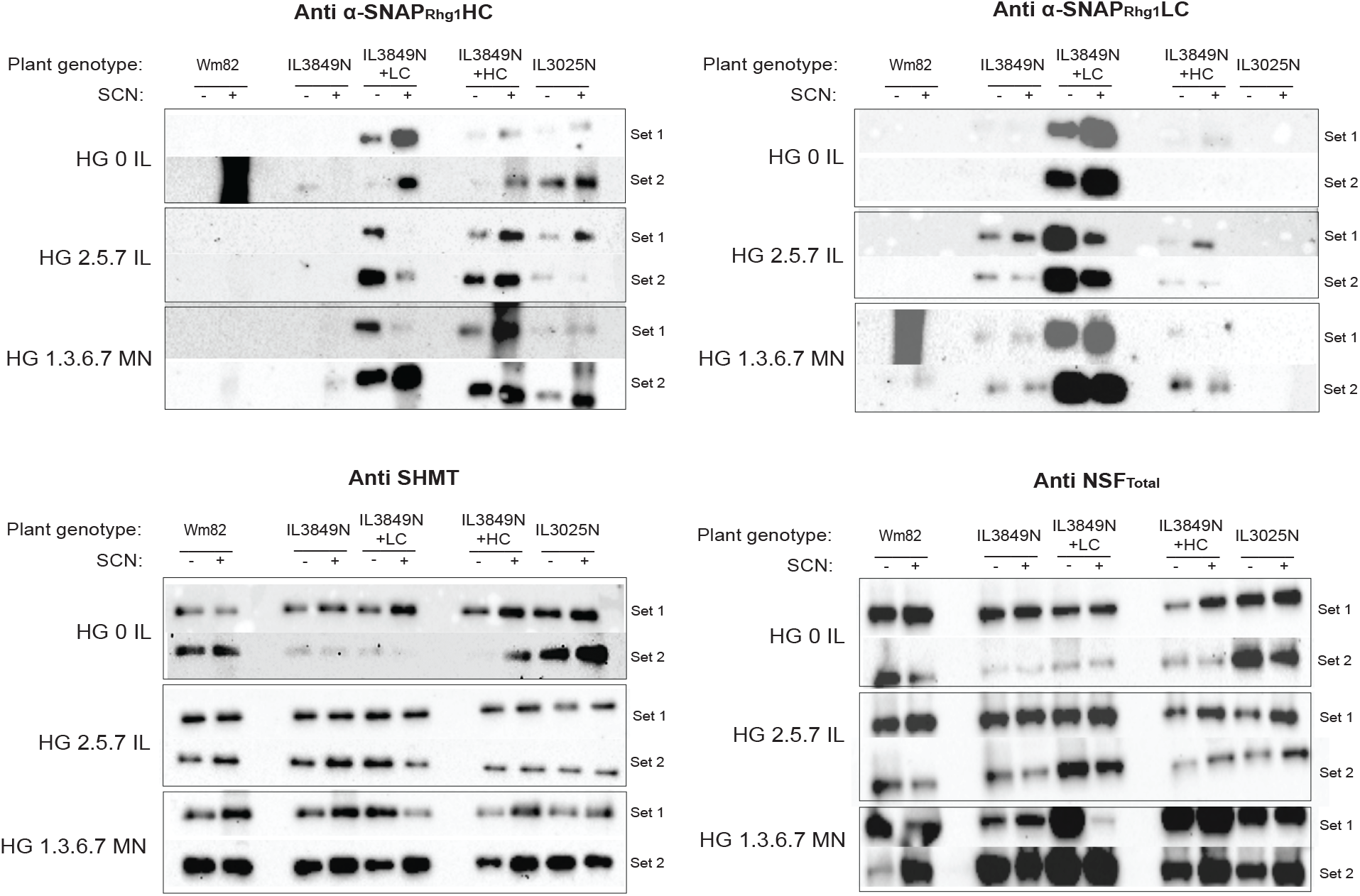
Protein abundance immunoblots for 7 dpi samples of SCN-infested root zones or similar areas from non-infested samples of Wm82 (Susceptible), IL3849N (*rhg1-a* parent), IL3849N+LC, IL3849N+HC and IL3025N (*rhg1-b* parent), using the indicated affinity-purified polyclonal antibodies. Antibody raised against α-SNAP_Rhg1_HC-specific C-terminus peptide cross-reacts with the α-SNAP_Rhg1_LC protein but not the Wm82 α-SNAP_Rhg1_WT, while antibody against α-SNAP_Rhg1_LC-specific C-terminus peptide is specific only to the α-SNAP_Rhg1_LC protein. All experiments were done on pooled sample of three roots each, separately for Set 1 and Set 2 seedlings. Equivalency of protein loading confirmed by Ponceau staining (Figure S4).

The abundance of transcripts encoding α-SNAP_Rhg1_HC exhibited different behaviors in response to different SCN populations (Figure 2). When comparing SCN-infested root zones (colored bars) to similar areas from non-infested samples (grey bars) at 7 dpi, significant or near-significant increases were observed in response to HG 2.5.7 IL while decreases were observed in response to HG 0 IL, along with a decrease in one of two lines in response to HG 1.3.6.7 MN. The α-SNAP_Rhg1_HC transgene in IL3849N+HC is expressed from a *GmUbi* (soybean ubiquitin) promoter rather than the IL3025N native *Glyma*.*18G022500* promoter, suggesting possible regulation by SCN that is specific to the mRNA sequence rather than the promoter.

There were no significant differences between plant genotypes in their fold-change of transcript abundance for the transcripts encoding α-SNAP_Rhg1_HC, NSF_RAN07_ and SHMT_*Rhg4*_ after challenge with the SCN populations HG 0 IL or HG 1.3.6.7 MN (Figure 2). With one exception, the same was true for α-SNAP_Rhg1_LC. However, for HG 2.5.7 IL, the fold change increase in SHMT_*Rhg4*_ transcript abundance after infection was less in most of the resistant lines compared to elevation of SHMT_*Rhg4*_ transcript in the SCN-susceptible Wm82 control (Figure 2, colored lines for p-value comparisons).

Polyclonal antibodies were used to monitor protein abundance at 7dpi for α-SNAP_Rhg1_LC, α-SNAP_Rhg1_HC, NSF_Total_ or SHMT_*Rhg4*_ with and without SCN infection (Figure 3, see also Figure S4 for Ponceau stain validation of equivalent loading between lanes). In these experiments the SCN population HG 2.5.7 IL elicited a different α-SNAP_Rhg1_LC protein abundance response than the HG 0 IL or HG 1.3.6.7 MN populations. Previous immunogold electron microscopy studies have documented ∼15-fold elevated abundance of α-SNAP_Rhg1_HC and α-SNAP_Rhg1_LC in syncytial cells (Bayless et al., 2016; Bayless et al., 2019). In the present study the SCN population HG 0 IL caused the expected increase in α-SNAP_Rhg1_LC abundance in the IL3849N+LC line. However, the abundance of α-SNAP_Rhg1_LC protein decreased in the IL3849N+LC lines during infection by the SCN population HG 2.5.7 IL (Figure 3). The same result was observed in additional experiments. This decrease in α-SNAP_Rhg1_LC protein abundance was not consistently observed in the parental IL3849N line that carries only the native *rhg1-a* locus. The HG 1.3.6.7 MN SCN elicited neither a decrease nor an increase in α-SNAP_Rhg1_LC abundance. This reproducible decrease in α-SNAP_Rhg1_LC protein abundance in response to HG 2.5.7 IL SCN was observed in two independent transgenic IL3849N+LC lines.

As might be predicted from previous studies (Bayless et al., 2016), in the native *rhg1-b* line IL3025N the abundance of α-SNAP_Rhg1_HC protein increased after inoculation for all tested SCN populations (5 of the 6 samples in Figure 3). Intriguingly, in IL3849N+HC (where α-SNAP_Rhg1_HC is expressed from a *GmUbi* promoter), the abundance of α-SNAP_Rhg1_HC protein also went up after inoculation for all tested SCN populations (6 of 6 samples in Figure 3). We parenthetically note that as previously documented, the custom anti-α-SNAP_Rhg1_HC polyclonal antibody weakly cross-reacts with α-SNAP_Rhg1_LC (Bayless et al., 2016). There is no α-SNAP_Rhg1_LC protein in *rhg1-b* IL3025N so the band detected by anti-α-SNAP_Rhg1_HC should entirely represent α-SNAP_Rhg1_HC abundance (Figure 3). The extent of background detection of α-SNAP_Rhg1_LC in the IL3849N+HC line probed with anti-α-SNAP_Rhg1_HC can be judged from the signal in the “IL3849N” lanes, as all lanes in the same immunoblot (same row within the composite Figure 3) were loaded with equivalent total protein. For the IL3849N+LC lanes in the blots probed with anti-α-SNAP_Rhg1_HC, we infer that the signal is from cross-detection of overexpressed α-SNAP_Rhg1_LC.

For SHMT_*Rhg4*_ and NSF_Total_ protein, there were few reproducibly significant protein abundance changes. We did, however, recurrently note a subtle increase in SHMT_*Rhg4*_ protein abundance in IL3849N+HC after inoculation with any of the SCN populations (Figure 3). NSF_Total_ also exhibited a subtle trend of increased abundance after HG 2.5.7 IL SCN infection in IL3849N+HC as well as in IL3025N (Figure 3). Table S2 provides a summary of the transcript and protein abundance findings from this study.

### Salicylic acid-responsive *NIMIN1* transcript, but not jasmonic acid-responsive *AAT* transcript, was upregulated in response to all SCN HG types in all tested soybean genotypes

In addition to genes directly associated with SCN resistance, we also monitored at 7dpi the transcript abundance of the JA pathway indicator gene AAT and the SA pathway indicator gene *NIMIN1* (Beyer et al., 2021). For inoculations with all three HG types, the abundance of AAT remained relatively unchanged from the basal (non-inoculated) level across genotypes, except in SCN-susceptible Wm82 (Figure S3). In contrast, the relative transcript abundance of SA pathway activity indicator gene (*NIMIN1*) went up at least 3-5 fold after SCN infection, and often more, for all of the tested plant genotypes and nematode HG types (Figure S3). Most of those increases after SCN inoculation were statistically significant (Figure S3). An ANOVA Tukey HSD test did not reveal any significant differences in the magnitude of the fold-change increase in *NIMIN1* across the plant genotypes or nematode treatments.

## (iv) Discussion

The present study documented incremental but significant improvements of SCN resistance in soybean, indicating that elevated expression of α-SNAP_Rhg1_ and/or combined expression of α-SNAP_Rhg1_LC and α-SNAP_Rhg1_HC protein types may be useful tools for improved control of SCN. Elevation of α-SNAP_Rhg1_ beyond what is provided by endogenous *rhg1-a* or *rhg1-b* haplotypes has not been achievable by conventional plant breeding. We elevated α-SNAP_Rhg1_ protein levels by transgenic methods, which was the most accessible approach. In the future such a trait alternatively might be created by mutagenesis, gene editing or discovery of soybean lines with increased copy number at the *Rhg1* locus – all of which can bypass expensive GMO plant deregulation processes. Transgenic elevation or diversification of α-SNAP_Rhg1_ might optimally be carried out using a fully cisgenic DNA construct that delivers native soybean DNA sequences, which would also limit GMO plant deregulation requirements. The choice of promoter will have a substantial impact on the resulting efficacy of SCN resistance.

It also has not been feasible via conventional plant breeding to create an individual soybean line that expresses both the α-SNAP_Rhg1_LC and α-SNAP_Rhg1_HC protein types, because the corresponding genes are allelic and soybean is an inbred crop that becomes homozygous at most loci. We and others speculate that some SCN of HG type 1.3.6 partially defeat *rhg1-a*/*Rhg4* but not *rhg1-b*, while some SCN of HG type 2.5.7 partially defeat *rhg1-b* but not *rhg1-a*/*Rhg4*, at least in part due to the different α-SNAP_Rhg1_ isoforms present in those lines. In the present work, presence of both α-SNAP_Rhg1_LC and α-SNAP_Rhg1_HC in a single IL3849N+HC soybean line enhanced *rhg1-a*/*Rhg4* SCN resistance and may improve the durability of the deployed SCN resistance over multiple seasons. An *Rhg1* haplotype that carries both *rhg1-a* and *rhg1-b* segments among its multiple ∼30kb *Rhg1* repeats in theory seems possible to generate and retain through conventional plant breeding, but to our knowledge has not been observed. Emergence of single heterologous *rhg1-a*/*rhg1-b* haplotypes may be suppressed due to poor meiotic pairing around *rhg1-a* and *rhg1-b*, as those existing haplotypes are typically ∼108kb and ∼312kb respectively (Bayless et al., 2019; Cook et al., 2014; Cook et al., 2012; Lee et al., 2015). Heterologous *rhg1-a*/*rhg1-b* presence within the same complex locus may alternatively be suppressed by gene conversion mechanisms, as there is evidence for ongoing homogenization of Rhg1 repeats (the sequences of the individual repeats are strikingly close to identical, Cook et al., 2014). Genetic transformation is a viable method to obtain single plant lines that express both α-SNAP_Rhg1_LC and α-SNAP_Rhg1_HC.

Overexpressing either α-SNAP_Rhg1_HC or α-SNAP_Rhg1_LC in IL3849N (*rhg1-a*/*Rhg4*) enhanced resistance against HG 0 and HG 2.5.7 SCN populations against which IL3849N already displays good resistance. It was noteworthy that by two metrics, cyst count at 30 dpi and post-harvest egg count in the soil, the transgenic lines exhibited enhanced resistance to SCN HG 2.5.7 IL (Figure 1 and Figure S2d). In the face of the ongoing evolution of SCN populations, resistance that reduces the end-of-season SCN egg count in the soil can be valuable both for long-term management of SCN disease pressure and for SCN resistance durability in individual fields. Stacking cisgenic elevation of α-SNAP_Rhg1_HC or α-SNAP_Rhg1_LC with other SCN resistance QTL that also incrementally restrict the success of SCN may further improve resistance durability, especially if those other QTL do not function through mechanisms related to α-SNAP proteins.

It was disappointing that overexpressing either α-SNAP_Rhg1_HC or α-SNAP_Rhg1_LC in a *rhg1-a*/*Rhg4* genetic background failed to significantly enhance resistance against an HG 1.3.6.7 SCN population. A possible interpretation of this result is that the HG 1.3.6.7 MN SCN population that was used has successfully overcome the resistance mechanism(s) mediated by any type of α-SNAP_Rhg1_. HG 1.3.6 SCN have partially overcome *rhg1-a*/*Rhg4* resistance but have not overcome the resistance conferred by *rhg1-b*, which expresses α-SNAP_Rhg1_HC but whose ∼10 *Rhg1* copies also more strongly express the resistance-conferring AAT_Rhg1_ and WI12_Rhg1_ proteins (Cook et al., 2014). This suggests manipulation of AAT_Rhg1_ and/or WI12_Rhg1_ proteins as an alternative means to achieve improved resistance against at least some subset of HG 1.3.6 SCN.

Many attempted transgenic disease resistance implementations are rejected because they cause a yield penalty (Collinge et al., 2010; van Esse et al., 2020). A modest growth/defense tradeoff is acceptable only in cases where disease pressure is sufficient to cause even greater loss of plant yield. That may be the case with SCN, meaning that a small compromise in yield potential in the absence of SCN may still be acceptable if it provides for elevated yields in the presence of SCN infestations, which are increasingly ubiquitous. A full-scale set of field-based yield studies may reveal some yield penalty from elevated expression of α-SNAP_Rhg1_, but the present results are encouraging in suggesting that such a penalty, if any, may be relatively minor. Modest yield penalties often can be overcome by further transgene engineering or by improvement of compatibility with other components of the plant genetic background. It was also encouraging that no changes in overall soybean seed protein, oil or fiber quantity were detected due to overexpression of α-SNAP_Rhg1_ in this study.

Overexpressing α-SNAP_Rhg1_HC in the high-copy *rhg1-b* IL3025N background did not detectably improve resistance against any of the three tested SCN HG types. It is possible that this constitutive overexpression did not significantly add appropriately localized SCN-responsive elevation of α-SNAP_Rhg1_HC to a line that already carries eight or nine copies of the endogenous gene. We also did not obtain any transformants in the *rhg1-b* background using a α-SNAP_Rhg1_LC construct. The absence of a resistance-type SHMT_*Rhg4*_ might be one plausible reason for this outcome (Lakhssassi, Piya, Knizia, et al., 2020).

When transcript and/or protein abundances were monitored for α-SNAP and a small set of other relevant loci, surprising differences in α-SNAP protein abundance were discovered. Previous work with HG 0 SCN has shown 10-20 fold upregulation of α-SNAP_Rhg1_HC or α-SNAP_Rhg1_LC protein abundance in syncytia (Bayless et al., 2016; Bayless et al., 2019). We again observed increased abundance of α-SNAP_Rhg1_HC or α-SNAP_Rhg1_LC protein in SCN-infested root segments for HG 0 SCN. However, we repeatedly observed a decrease in α-SNAP_Rhg1_LC protein abundance in samples from two independent IL3849N+LC lines after inoculation with the HG 2.5.7 IL SCN population. The lines exhibited good SCN resistance, but a decrease in α-SNAP_Rhg1_LC protein abundance compared non-inoculated plants of the same genotype. Abundance of the transcript encoding α-SNAP_Rhg1_LC was not decreased by inoculation with HG 2.5.7 IL SCN. The decrease in α-SNAP_Rhg1_LC protein abundance was not observed with the HG 0 or HG 1.3.6.7 SCN populations used in this study. Furthermore, the HG 2.5.7 IL SCN population did not reproducibly decrease the already-lower α-SNAP_Rhg1_LC abundance present in IL3849N lacking transgenically ovexpressed α-SNAP_Rhg1_LC. The HG 2.5.7 IL SCN population also did not detectably decrease α-SNAP_Rhg1_HC or α-SNAP_Rhg1_LC protein abundance in the IL3849N+HC line. The lower α-SNAP_Rhg1_LC protein abundance after infestation by HG 2.5.7 IL SCN was still at or above levels of the α-SNAP_Rhg1_LC protein in the parent IL3849N line in the same experiment, but the induced decrease is intriguing. The full set of results seem to suggest that the HG 2.5.7 IL SCN population we used does not carry some capacity to cause specific degradation of α-SNAP_Rhg1_LC protein or to reduce translation of its transcript. We instead speculate that the overabundance of α-SNAP_Rhg1_LC protein in the IL3849N + LC lines may alter α-SNAP/NSF/SNARE interactions or vesicle trafficking that is also impacted by the HG 2.5.7 IL SCN (but not the HG 0 or HG 1.3.6.7 SCN used in this study), leading to the observed strong decrease in α-SNAP_Rhg1_LC protein abundance.

We noted a separate transcriptional phenomenon with the SCN population HG 2.5.7 IL. The abundance of transcripts encoding α-SNAP_Rhg1_HC (not LC) increased in response to HG 2.5.7 IL both in the native context of IL3025N (p = 0.050; carries the PI 88788-type *rhg1-b* locus) and in the IL3849N+HC line (p = 0.133; carries the Peking-type *rhg1-a* locus and a transgene overexpressing α-SNAP_Rhg1_HC). The transgene is expressed from a *GmUbi* (soybean ubiquitin) promoter while the IL3025N version is from the native *Glyma*.*18G022500* (*GmSNAP18*) gene, suggesting possible transcript abundance regulation that is specific to the mRNA sequence rather than the promoter. Under the same experimental conditions, HG 0 IL SCN caused the opposite transcript response and decreased the abundance of the transcript encoding α-SNAP_Rhg1_HC in both IL3025N and the IL3849N+HC line (p = 0.032 and 0.068; Figure 2), as did HG 1.3.6.7 MN nematodes on the IL3849N+HC line (p = 0.012). The observations of this and the preceding paragraph open opportunities for further study of HG 2.5.7 SCN parasitism that are beyond the scope of the present report.

We did not detect other strongly surprising results in the transcript and protein abundance experiments of this study, but a few additional observations are worthy of note. Although α-SNAP_Rhg1_LC and SHMT_*Rhg4*_ have been shown to physically interact and are necessary partners for plant viability and resistance (Lakhssassi, Piya, Bekal, et al., 2020; Lakhssassi, Piya, Knizia, et al., 2020), neither overexpressing α-SNAP_Rhg1_LC nor SCN challenge detectably altered SHMT_Rhg4_ transcript or protein abundance in most of the experiments conducted here. However, in only the SCN-susceptible Wm82 accession, SCN population HG 2.5.7 IL did reproducibly elevate the abundance of the transcript encoding SHMT_Rhg4_. Separately, in 14 of the 15 interactions tested (the exception being Wm82 inoculated with HG 0 SCN), no increase in transcript abundance for the JA pathway indicator gene AAT was observed, while substantial increases in the SA pathway indicator gene *NIMIN1* were observed in all 15 of the tested interactions, mirroring the findings from at least one other study (Kandoth et al., 2011).

Returning to the primary focal point of the present study, elevated expression of α-SNAP_Rhg1_ proteins incrementally but reproducibly improved the SCN resistance of soybean lines against SCN populations toward which the parental line already exhibited moderate or strong resistance. There are extensive opportunities for further refinement and testing of these technologies to improve SCN resistance in soybean.

## (v) Experimental procedures

### Soybean Transformation

Transgenic soybean lines were generated by the University of Wisconsin – Madison Wisconsin Crop Innovation Center using their standardized meristem-based soybean transformation method, a hormone-free method for *Agrobacterium tumefaciens*-mediated transformation of embryonic plantlets extracted from imbibed mature seeds. Seeds of SCN-susceptible soybean variety *Williams 82* and the high-yielding SCN-resistant public varieties IL3849N (LD07-3395bf) and IL3025N (LD11-2170) were genetically transformed to produce the lines used in the study. IL3849N is homozygous for *rhg1-a* as well as for the resistance-associated allele at *Rhg4* but not rhg2. IL3025N is homozygous for *rhg1-b*. Overexpression of soybean *Glyma*.*18G022500* (*GmSNAP18*) coding sequences (CDS) encoding either α-SNAP_Rhg1_LC (Low copy type or *rhg1-a* type) or α-SNAP_Rhg1_HC (High copy type or *rhg1-b* type) under control of a 917bp soybean ubiquitin promoter (*GmUbi*; Genbank submission U310508) (Concibido et al., 1997; Hernandez-Garcia et al., 2009) was achieved with gene cassettes that also used an *Agrobacterium nos terminator*. The adjacent spectinomycin selection marker (*aadA* product) was fused to petunia chloroplast-targeting peptide and driven by a CaMV 35S promoter.

### Soybean line selection

Transgenic lines were chosen for study based on elevated production of α-SNAP_Rhg1_ protein in pooled root samples from >12 sibling seedlings, determined by protein immunoblots probed with affinity purified polyclonal 1° antibody raised against either “Ac-C(dPEG4)EQHEAIT-OH” peptide for α-SNAP_Rhg1_HC or “Ac-C(dPEG4)EEYEVIT-OH” peptide for α-SNAP_Rhg1_LC (Bayless et al., 2016), as described below and shown in Figure 3. Lines for further analysis were identified based on uniform spectinomycin resistance across T2 plants, assessed 7 days after leaflets were sprayed with 0.1% (m/v) spectinomycin in 0.1% (v/v) Tween 20 surfactant. Lines were further chosen based on apparent homozygous single-copy transgene copy number relative to the endogenous single-copy gene Glyma.18G022800, assessed in T2 plants by qPCR using CTAB-extracted genomic DNA as template, with oligonucleotide primers p23SF: CTAGCCAATACGCAAACCGC and p23SR: CCTGGGGTGCCTAATGAGTG and reference gene primers 2620F: AAGCCCAACAGGCCAAAGAGAG and 2620R: ACACCAAATGGGTTCGCACTTC. Evaluated T2 and T3 seed lines were used for further studies. The acceptable lines were from two separate transformation events for α-SNAP_Rhg1_LC OE in IL3849N (212-7 and 212-1) but only one successful event was obtained for α-SNAP_Rhg1_HC OE in IL3849N (213-1), after which separate lines from two different 213-1 T1 progeny were established and utilized.

### SCN resistance tests

SCN reproduction (cyst formation) relative to an SCN-susceptible control line was quantified on seedlings grown in a controlled environment growth room. The seeds were organized into two sets for the biological replicate experiments: Set1 utilized 212-7 and 213-1-1 for α-SNAP_Rhg1_LC and α-SNAP_Rhg1_HC overexpression respectively while Set2 utilized 212-1 and 213-1-3. Twelve plants of each genotype were planted in a completely randomized and blinded design in a heat-treated 50:50 sand:clay soil mixture and were inoculated with 1,000 SCN eggs of the chosen HG type. Plants were incubated in a growth room under 14h daylight at 27 °C day and 22 °C night temperatures. All cysts were extracted from each cup at 35 dpi and enumerated. The female index was calculated using the formula: (number of cysts on test line/average number of cysts on Williams 82 plants) x 100.

### Nematodes

Inbred soybean cyst nematode populations of HG type 0 and HG type 2.5.7 were obtained from Alison Colgrove at University of Illinois, Urbana-Champaign and were maintained on Williams 82 and IL3025N respectively. An inbred HG 1.3.6.7 population was obtained from Senyu Chen at University of Minnesota and maintained subsequently on IL3849N. These populations are referred to as HG 0 IL, HG 2.5.7 IL and HG 1.3.6.7 MN. Freshly hatched second-stage juveniles of all HG type populations used in this study produced similar numbers of cysts on the susceptible Wm82 (Figure S1a).

### Greenhouse Yield Assay

All soybean seed yield assays were performed in the same University of Wisconsin – Madison greenhouse using a randomized complete block design with a border of IL3849N plants surrounding the four blocks, and blocks arranged for within-block environmental uniformity. The greenhouse was maintained at no less than 27°C day and 22°C night temperatures. For winter experiments, illumination was 16h until 60 dpi, then 14 h until 100 dpi, then 12h until dry seed harvest. To initiate the experiment, seeds were germinated on paper towels until they developed 5-7 cm radicle and then three seedlings of the same genotype were planted in a 20 L plastic pot with 50:50 mixture of untreated fine sand and clay soil, with or without SCN HG 2.5.7 IL. For treatments with SCN HG 2.5.7 IL, 300,000 SCN eggs were mixed into the top 10L of sand:soil mixture before planting the seedlings. The experiment was conducted three times (Winter 2020: Set 1 seeds, Summer 2021: Set 1 seeds, and Winter 2021: Set 2 seeds). Pots were irrigated alternately with water or fertilizer water (300 ppm of nitrogen, 150 ppm of phosphorus, and 300 ppm of potassium) until maturity. Plants were left to dry without irrigation for a week post maturity preceding harvest. Pods were harvested, threshed, and weighed manually. Seed moisture, protein, oil, and fiber content were evaluated using a Perten Instruments IM9500 NIR spectroscope. For the winter experiments, the seeds from all four technical replicates from each treatment were pooled and three subsamples were used to calculate the average seed composition. For the summer experiment, seeds from each technical replicate of every treatment were analyzed separately and the average seed composition was computed.

### Field Yield Assay

Transgenic IL3849N lines and the parent IL3849N control were evaluated for yield in field conditions under USDA-APHIS-BRS permit at University of Nebraska–Lincoln field facilities located in Saunders County, NE, USA. Seed from the greenhouse yield assay was planted in a randomized complete block design on 24 May 2023 for IL3849N, two events of IL3849N+LC (212-7-2 and 212-1-1), and two lines from the single IL3849N+HC event (213-1-1 and 213-1-3). The four blocks each contained two adjacent 3m rows of each of the five lines and the entire test was bordered by IL3849N on all four sides. Prowl H_2_0 (at 4.2 L/ha), Sonalan (at 700 mL/ha) and Canopy XL (at 175 mL/ha) were sprayed the day after planting along with irrigation. The experiment plot had been planted to Bt/Roundup Ready corn in 2022, which had received Harness Extra (at 5.6 L/ha) and Atrazine (at 2.8 L/ha). The 2023 plots were overhead-irrigated a total of 6 times to supplement precipitation. Plots were harvested on 10 October 2023 and the seeds were weighed by each two-row plot. After harvest, six six-inch deep soil cores were sampled in a zig-zag manner and pooled for each two-row plot, and then sent for SCN testing at the University of Nebraska-Lincoln Extension SCN testing program.

### SCN-Induced Transcript Abundance Assay

Paper-germinated soybean seedlings of Wm82, IL3849N, IL3849N+LC and IL3849N+HC were planted into 5 cm cups in heat-treated fine sand and inoculated with 1,000 SCN J2s of either HG 0 IL, HG 2.5.7 IL, HG 1.3.6.7 MN, or no nematodes. The setup was irrigated with tap water and grown in a growth chamber with 14h daylight, and 27°C day and 22°C night temperatures. The experiment was performed twice, once with Set 1 seed and once with Set 2 seed. Each time a randomized complete block design was planted with six blocks (single plant per pot), three blocks of which were harvested for RNA work and three for protein work. At 7 dpi the roots were rinsed with room temperature tap water and lateral roots of SCN-inoculated plants were examined under a stereoscope for lesions or nematode entry points. Infested root areas were harvested with 1 cm margins on either side of the lesion(s). Multiple root sections from individual plants were combined into single ice-chilled 2mL cryovials with five 2mM ceramic beads each, and flash frozen in liquid nitrogen within three minutes of the initial tap water rinse of the plant roots. Root samples were stored at -80°C. Total RNA was then extracted using Zymo Direct-Zol RNA miniprep kit as per manufacturer’s instructions. Stock cDNA was synthesized from 1.5 μg of total RNA using Solis Biodyne FIREScript kit following the manufacturer’s protocol. cDNA quality was checked on a 2% agarose gel using the *Cyp2* primers (*Glyma*.*12G024700*) using an end-point PCR. Primer efficiencies of all oligonucleotide primers used (Table S3) were verified to be >97% using 4 serial dilutions of 10-fold each (Beyer et al., 2021). qRT-PCR reactions for two replicates each of two separate roots for each treatment were were performed using Solis Biodyne HOT FIREPol EvaGreen qPCR supermix in a BioRad CFX96 Realtime system as per manufacturer’s protocol. For every gene, mean Ct values for each root were calculated from the two technical replicates. For all four plants (two from each biological rep), the dCt values with respect to *NREG-2* reference gene were calculated. A ddCt value for each inoculated root with respect to the corresponding non-inoculated root of the same plant genotype on the same qPCR plate was calculated. The resulting graphs show the mean 2^(-ddCt) values (fold-change in transcript abundance) for four independent samples. All statistical analyses were performed in R and the data were visualized using Microsoft Excel.

### SCN-Induced Protein Expression Assay

For protein abundance determinations, SCN-infested root zones or similar areas from non-infested samples issue were obtained from other blocks within the same plantings as described above for transcript abundance. Root samples from three replicates of the same treatment were pooled and equally split into 3 subsamples of 50 mg each and flash frozen with liquid nitrogen in 2mL cryovials with 5 2mM ceramic beads. These samples were stored immediately at -80 °C until used. Protein from root samples was extracted and concentrations were normalized as for line selection protein immunoblots (see above). After SDS-PAGE samples were transferred to Nitrocellulose. Previously validated rabbit anti-α-SNAP_Rhg1_LC (Bayless et al., 2016), rabbit anti-α-SNAP_Rhg1_HC (Bayless et al., 2016) and rabbit anti-NSF (Bayless et al., 2016) (raised by Pacific Immunology against synthetic peptide representing residues 300 to 324 of soybean NSF_Chr07_ “ETEKNVRDLFADAEQDQRTRGDESD”; will recognize both soybean NSF proteins), and anti-SHMT (Serine Hydroxy Methyl Transferase; from Agrisera #AS05075) were used as per (Bayless et al., 2016). Detection utilized goat anti-rabbit peroxidase conjugated secondary antibody and Thermo Scientific SuperSignal^TM^ West Dura extended duration substrate. The blots were repeated with both Set1 and Set2 lines for each combination of antibody, plant genotype, nematode inoculation, and nematode HG type.

## Supporting information

Three supplemental tables and Four supplemental figures

## Author Contributions

DH co-conceived the experiments and conducted the growth-chamber, greenhouse, western blot assays and significantly contributed to the qPCR assays. DH also analyzed results and co-authored the manuscript. SD designed haplotype-specific qPCR assays and performed the qPCR assays. EN and RZ performed initial evaluation of the transgenic candidate lines and helped with growth-chamber assays. AFB co-conceived the experiments, co-analyzed the results, co-authored the manuscript, obtained project funding and served as group leader. All authors helped to revise the manuscript.

## (vi) Accession numbers

(not applicable)

## (vii) Acknowledgements

This project was supported by the United Soybean Board and the Wisconsin Soybean Marketing Board. We thank Tom Clemente and Derrick Grunwald for extensive contributions to preceding phases of this project and for helpful comments, Shaojie Han for construction of the transgene constructs, and WCIC staff members Rob Hornish and Mark Thompson for generation of the transgenic soybean lines. The authors are also grateful to Patrick Tenopir for facilitating field experiments, Adam Roth for assistance with the NIR seed analyses, Lynn Hummel, Ben Erdman, Deena Paterson, Issac Kabera, Ken Kmeicik, Jessica Peterson, Sydney Oerter, and John Stuntebeck for assistance with greenhouse operations, and multiple Bent lab members for constructive suggestions. The authors have no conflict of interest to declare; AB and RZ are inventors on patent applications related to *Rhg1* and NSF_RAN07_ that are owned by the Wisconsin Alumni Research Foundation.

## (viii) Short legends for Supporting Information

**Figure S1**. Virulence of the SCN populations on a susceptible host and schematic overview of the study.

**Figure S2**. Additional SCN reproduction and seed yield data.

**Figure S3**. Additional transcript abundance data and additional statistical tests for transcript data.

**Figure S4**. Ponceau stain images of the blots used in Figure 3.

## (x) Tables

(none; see Supplemental Tables)

## (Xi) Figure legends

**Figure S1**. Virulence of the SCN populations on a susceptible host and schematic overview of the study. (a) SCN cyst numbers for HG type populations used in the study on susceptible Wm82. (b) Schematic overview of this study. The error bars represent standard errors.

**Figure S2**. Additional SCN reproduction and seed yield data. Female indices (FI) on parent IL3025N *rhg1*-b line compared to IL3025N+HC for (a) SCN HG 0 IL (b) SCN HG 2.5.7 IL and (c) SCN HG 1.3.6.7 MN. FI experiments for SCN HG 0 IL and SCN HG 2.5.7 IL were done twice with 12 replicates each in a randomized design. FI experiment for SCN HG 1.3.6.7 MN were performed once with 15 replicates. (d) Greenhouse end-of-season SCN egg density (eggs/100 cc soil) comparisons for the parent IL3849N rhg1-a line, IL3849N+LC and IL3849N+HC with respect to the susceptible Wm82. (e) Field yield in g/plot for one experiment with IL3849N, IL3849N+LC (2 events) and IL3849N+HC (1 event, 2 lines) with 4 blocked replicates. The error bars represent standard errors. Only significant Tukey HSD p-values are shown.

**Figure S3**. Additional transcript abundance data and additional statistical tests for transcript data. (a) Transcript abundance (qPCR) comparisons for JA and SA defense pathway indicator genes. Tissue samples taken seven days after inoculation with the indicated SCN population or at same time point for non-inoculated control plants, for soybean roots of Wm82 (Susceptible), IL3849N (*rhg1-a* parent), IL3849N+LC, IL3849N+HC and IL3025N (*rhg1-b* parent). Statistical p-values are for comparisons of non-inoculated (grey bars) and SCN-inoculated (colored bars) samples of the same plant genotype, using two-tailed t-tests. (b) Additional ANOVA and Tukey HSD tests for the data of Figure 2 and Figure S3a, for comparisons of same plant genotype inoculated with different SCN HG types. For both (a) and (b), transcript fold-change (y-axis) is relative to non-inoculated roots of the same genotype. Error bars show standard error of the mean. All experiments were done on two roots each from Set 1 and Set 2 seeds. Each root had two qPCR technical replicates.

**Figure S4**. Ponceau stain images of the blots used in Figure 3. Total protein concentration was determined by Bradford assay for an aliquot from each root protein sample. Samples were then normalized within an experiment (horizontal set of lanes from the same blot) so that similar amounts of total protein were loaded into each lane.

