## Supplementary material for "Overexpression of α-SNAP_Rhg1_ can improve *rhg1-a* mediated soybean resistance to soybean cyst nematode": Three supplemental tables and Four supplemental figures

(a)

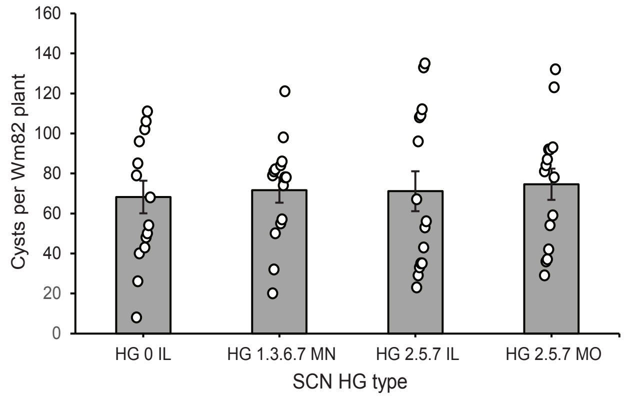

(b)

| Agrobacterium mediated transformation with Spectinomycin selection done at WCIC | | | |
|---|---|---|---|
| GmUbi OE<br>NOS Terminator<br>Background | +Empty vector (EV) | +HC α-SNAPRhg1 | +LC α-SNAPRhg1 |
| IL3849N (LC/ *rhg1*-a type) | No events | 213-1 (one event) | 212-7, 212-1<br>(two events) |
| IL3025N (HC/ *rhg1*-b type) | 202-3 (one event) | 205-1 (one event) | No events |

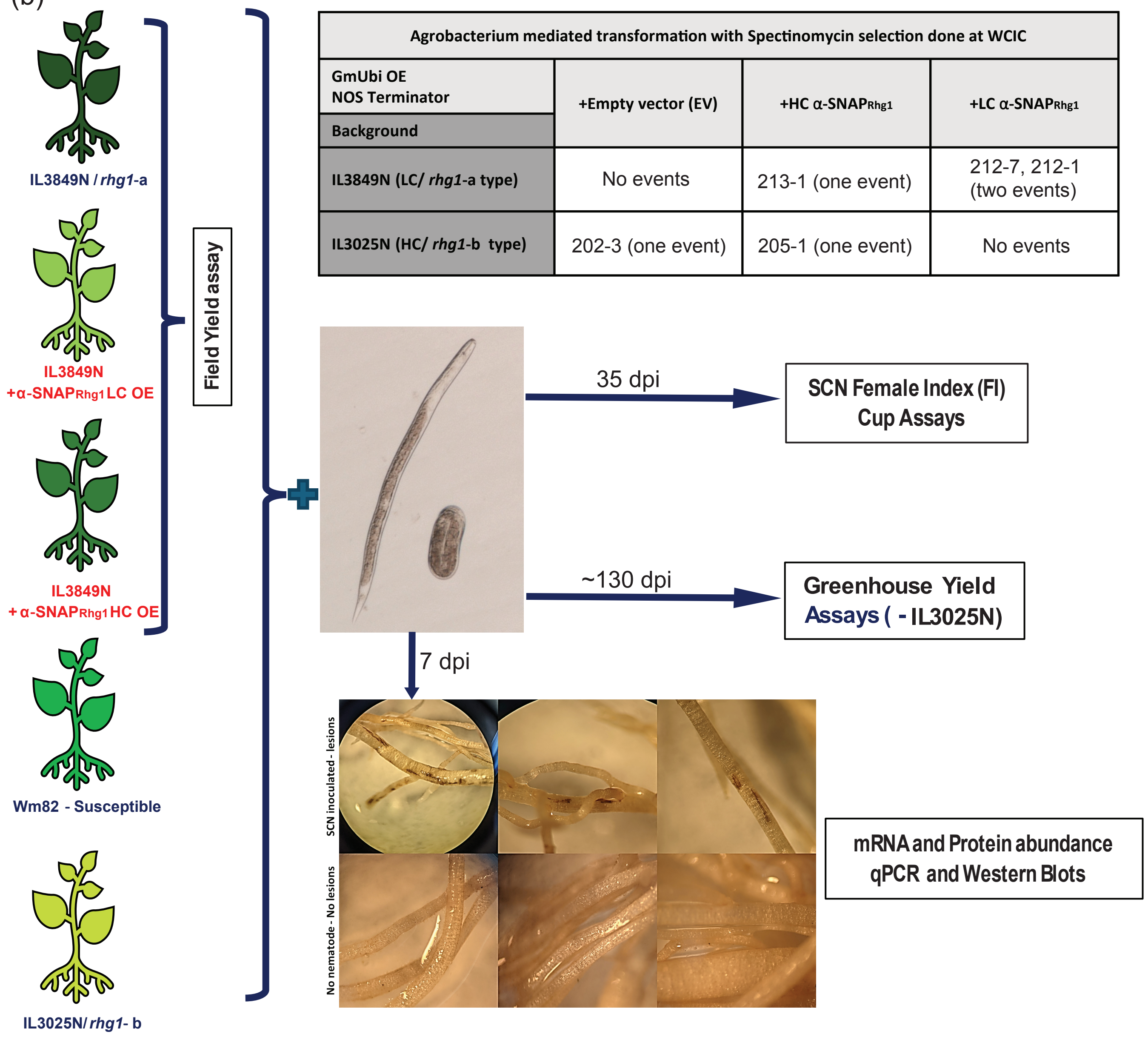

**Figure S1.** Virulence of the SCN populations on a susceptible host and schematic overview of the study. (a) SCN cyst numbers for HG type populations used in the study on susceptible Wm82. (b) schematic overview of this study. The error bars represent standard errors.

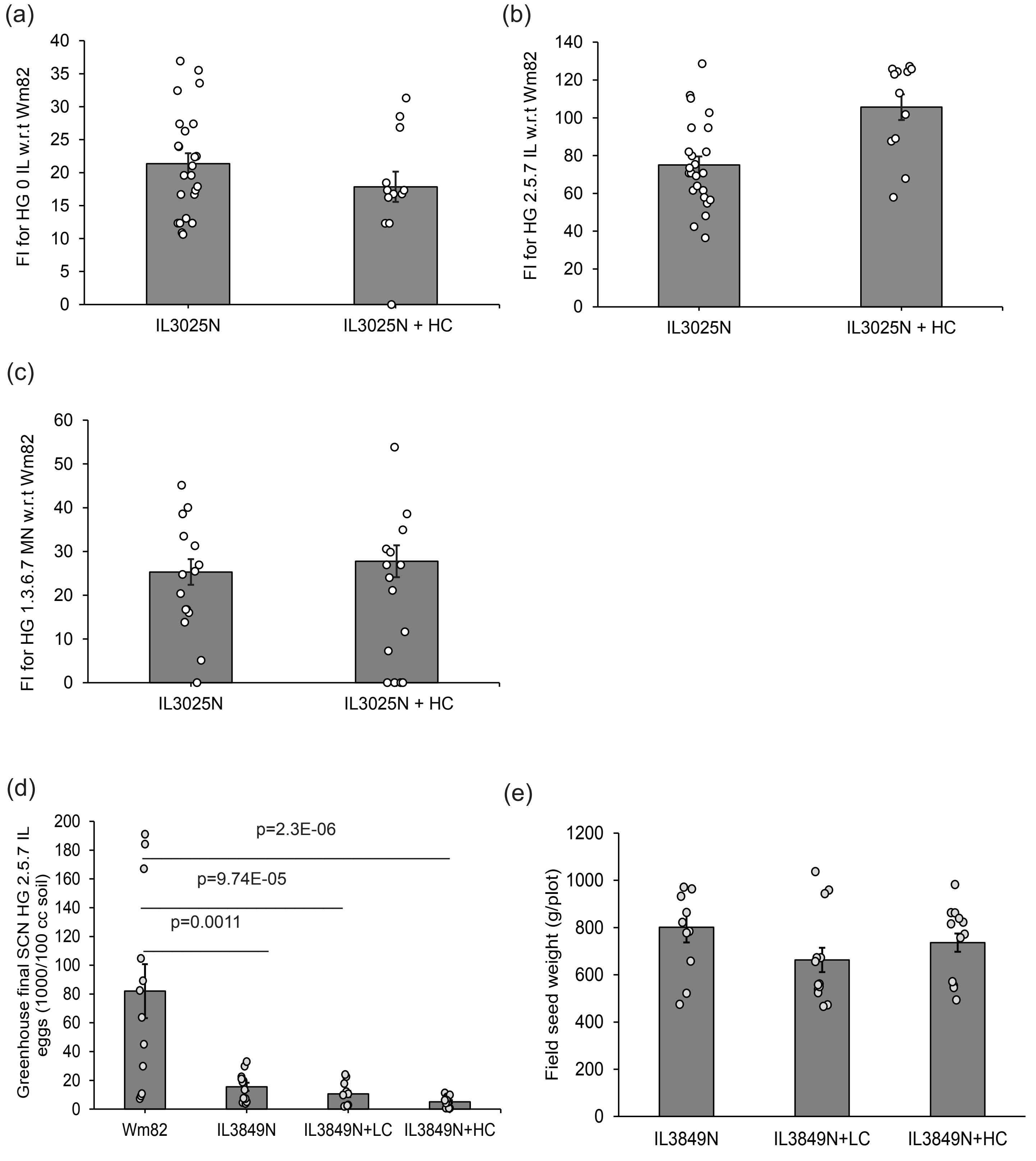

**Figure S2.** Additional SCN reproduction and seed yield data. Female indices (FI) on parent IL3025N *rhg1-b* line compared to IL3025N+HC for (a) SCN HG 0 IL (b) SCN HG 2.5.7 IL and (c) SCN HG 1.3.6.7 MN. FI experiments for SCN HG 0 IL and SCN HG 2.5.7 IL were done twice with 12 replicates each in a randomized design. FI experiment for SCN HG 1.3.6.7 MN were performed once with 15 replicates. (d) Greenhouse end-of-season SCN egg density (eggs/100 cc soil) comparisons for the parent IL3849N *rhg1-a* line, IL3849N+LC and IL3849N+HC with respect to the susceptible Wm82. (e) Field yield in g/plot for one experiment with IL3849N, IL3849N+LC (2 events) and IL3849N+HC (1 event, 2 lines) with 4 blocked replicates. The error bars represent standard errors. Only significant Tukey HSD p-values are shown.

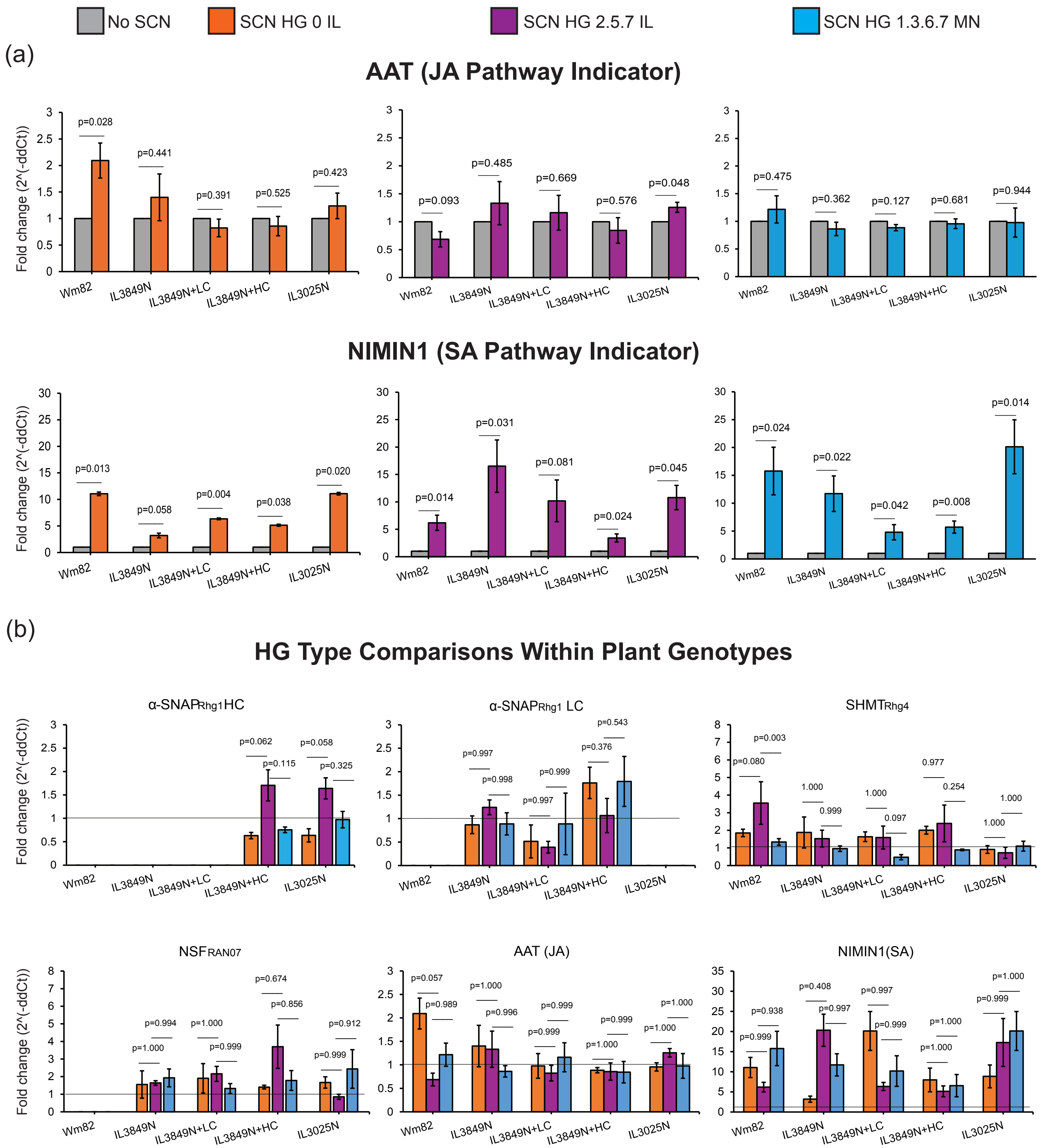

**Figure S3.** Additional transcript abundance data and additional statistical tests for transcript data. (a) Transcript abundance (qPCR) comparisons for JA and SA defense pathway indicator genes. Tissue samples taken seven days after inoculation with the indicated SCN population or at same time point for non-inoculated control plants, for soybean roots of Wm82 (Susceptible), IL3849N (*rhg1-a* parent), IL3849N+LC, IL3849N+HC and IL3025N (*rhg1-b* parent). Statistical p-values are for comparisons of non-inoculated (grey bars) and SCN-inoculated (colored bars) samples of the same plant genotype, using two-tailed t-tests. (b) Additional ANOVA and Tukey HSD tests for the data of Figure 2 and Figure S3a, for comparisons of same plant genotype inoculated with different SCN HG types. For both (a) and (b), transcript fold-change (y-axis) is relative to non-inoculated roots of the same genotype. Error bars show standard error of the mean. All experiments were done on two roots each from Set 1 and Set 2 seeds. Each root had two qPCR technical replicates.

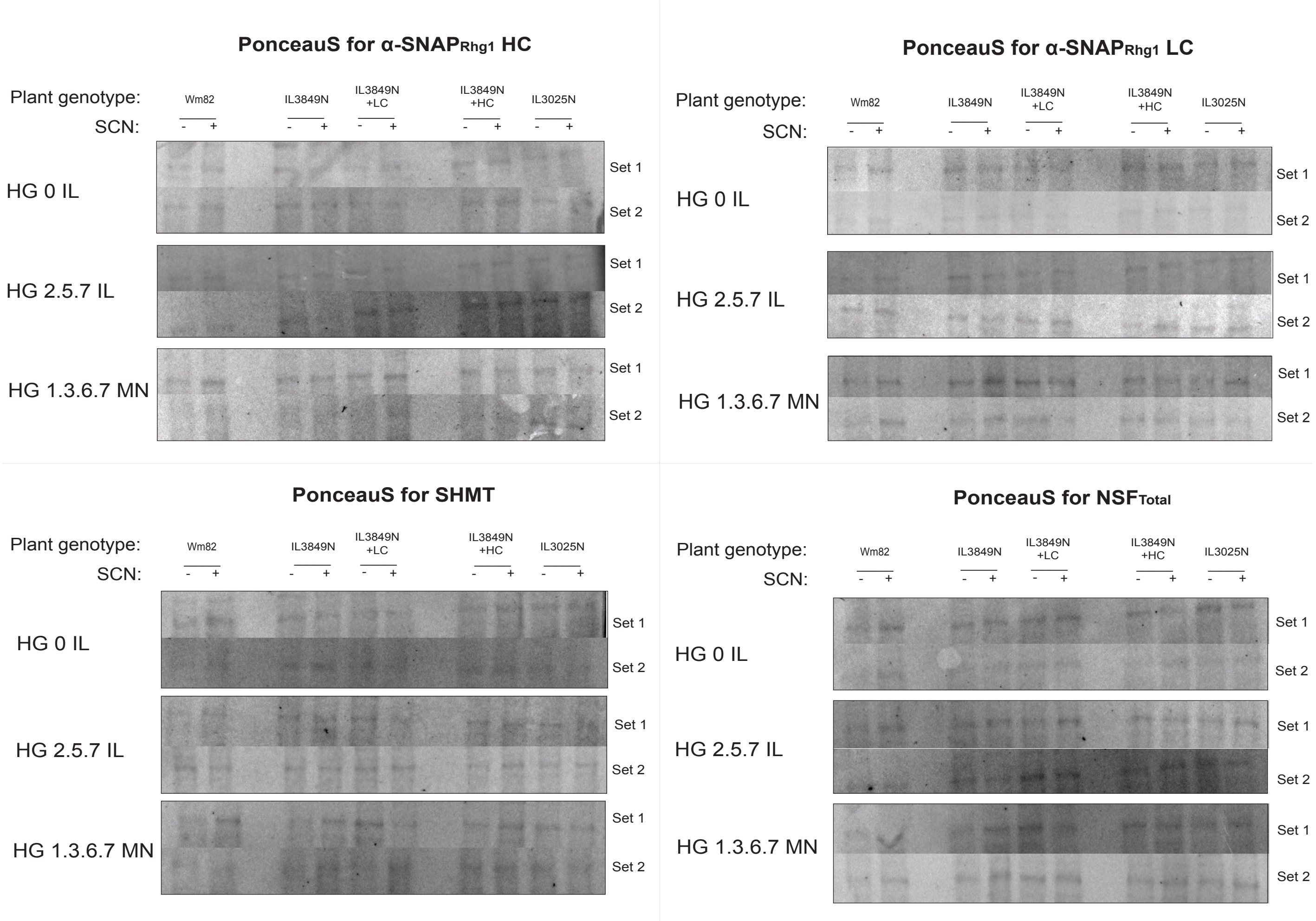

**Figure S4.** Ponceau stain images of the blots used in Figure 3. Total protein concentration was determined by Bradford assay for an aliquot from each root protein sample. Samples were then normalized within an experiment (horizontal set of lanes from the same blot) so that similar amounts of total protein were loaded into each lane.

**Supplemental Tables**

*Table S1. Greenhouse seed quality evaluation. ANOVA with only significant Tukey HSD p-values shown.*

| Signif. codes: 0 '***' 0.001 '**' 0.01 '*' 0.05 '.' 0.1 ' ' 1 | | Greenhouse seed oil content | | | | |
|---|---|---|---|---|---|---|
| **ANOVA** | | | | | | |
| | **Df** | **Sum Sq** | **Mean Sq** | **F value** | **Pr(>F)** | **Signif.** |
| Experiment | 2 | 12.51 | 6.2548 | 26.7662 | 2.67E-09 | *** |
| Nematode inoculation | 1 | 3.234 | 3.2339 | 13.8387 | 4.06E-04 | *** |
| Plant genotype | 3 | 53.51 | 17.8366 | 76.3282 | < 2.2e-16 | *** |
| Experiment x block | 9 | 3.054 | 0.3393 | 1.4521 | 0.183932 | |
| Nematode inoculation x Plant genotype | 3 | 0.998 | 0.3328 | 1.424 | 0.24336 | |
| Residuals | 68 | 15.89 | 0.2337 | | | |
| **Tukey HSD** | | | | | | |
| | | **Experiment** | **P.adj** | **Signif.** | | |
| | | Summer 2021 - Winter 2020 | 0 | *** | | |
| | | Winter 2021 - Winter 2020 | 0 | *** | | |
| | | Winter 2021 - Summer 2021 | 0.439032 | | | |
| | | **Nematode inoculation** | | | | |
| | | SCN - No SCN | 4.35E-04 | *** | | |
| | | **Plant genotype** | | | | |
| | | IL3849N+HC - IL3849N | 0.038878 | * | | |
| | | IL3849N+LC - IL3849N | 0.917721 | | | |
| | | Wm82 - IL3849N | 0 | *** | | |

| | | | | | | |
|---|---|---|---|---|---|---|
| | | IL3849N+LC - IL3849N+HC | 0.004996 | ** | | |
| | | Wm82 - IL3849N+HC | 0 | *** | | |
| | | Wm82 - IL3849N+LC | 0 | *** | | |
| | | **Nematode inoculation x Plant genotype** | | | | |
| | | **only significant interactions** | | | | |
| | | No SCN:Wm82-No SCN:IL3849N | 0 | *** | | |
| | | SCN:Wm82-No SCN:IL3849N | 1E-07 | *** | | |
| | | No SCN:Wm82-SCN:IL3849N | 0 | *** | | |
| | | SCN:Wm82-SCN:IL3849N | 2E-07 | *** | | |
| | | SCN:IL3849N+LC-No SCN:IL3849N+HC | 0.004086 | ** | | |
| | | No SCN:Wm82-No SCN:IL3849N+HC | 0 | *** | | |
| | | SCN:Wm82-No SCN:IL3849N+HC | 0.000342 | *** | | |
| | | No SCN:Wm82-SCN:IL3849N+HC | 0 | *** | | |
| | | SCN:Wm82-SCN:IL3849N+HC | 3.7E-06 | *** | | |
| | | No SCN:Wm82-No SCN:IL3849N+LC | 0 | *** | | |
| | | SCN:Wm82-No SCN:IL3849N+LC | 2E-07 | *** | | |
| | | No SCN:Wm82-SCN:IL3849N+LC | 0 | *** | | |
| | | SCN:Wm82-SCN:IL3849N+LC | 0 | *** | | |
| Signif. codes: 0 ‘***’ 0.001 ‘**’ 0.01 ‘*’ 0.05 ‘.’ 0.1 ‘ ’ 1 | | **Greenhouse seed protein content** | | | | |
| **ANOVA** | | | | | | |
| | **Df** | **Sum Sq** | **Mean Sq** | **F value** | **Pr(>F)** | **Signif.** |
| Experiment | 2 | 56.829 | 28.4147 | 6.8424 | 0.001961 | ** |
| Nematode inoculation | 1 | 1.671 | 1.6713 | 0.4025 | 5.28E-01 | |
| Plant genotype | 3 | 53.35 | 17.7832 | 4.2823 | 0.007907 | ** |
| Experiment x block | 9 | 8.538 | 0.9487 | 0.2284 | 0.989274 | |
| Nematode inoculation x Plant genotype | 3 | 5.222 | 1.7406 | 0.4191 | 0.739836 | |

| | | | | |
|---|---|---|---|---|
| Residuals | 68 | 282.385 | 4.1527 | |
| **Tukey HSD** | | | | |
| | | **Experiment** | **P.adj** | **Signif.** |
| | | Summer 2021 - Winter 2020 | 0.353527 | |
| | | Winter 2021 - Winter 2020 | 0.111191 | |
| | | Winter 2021 - Summer 2021 | 0.001335 | ** |
| | | **Nematode inoculation** | | |
| | | SCN - No SCN | 5.30E-01 | |
| | | **Plant genotype** | | |
| | | IL3849N+HC - IL3849N | 0.333203 | |
| | | IL3849N+LC - IL3849N | 0.991523 | |
| | | Wm82 - IL3849N | 0.034888 | * |
| | | IL3849N+LC - IL3849N+HC | 0.184874 | |
| | | Wm82 - IL3849N+HC | 0.621523 | |
| | | Wm82 - IL3849N+LC | 0.013905 | * |
| | | **Nematode inoculation x Plant genotype** | | |
| | | **only significant interactions** | | |
| | | NONE | | |

| Signif. codes: 0 ‘***’ 0.001 ‘**’ 0.01 ‘*’ 0.05 ‘.’ 0.1 ‘ ’ 1 |  | Greenhouse seed fiber content |  |  |  |  |
|---|---|---|---|---|---|---|
| **ANOVA** | | | | | | |
| | **Df** | **Sum Sq** | **Mean Sq** | **F value** | **Pr(>F)** | **Signif.** |
| Experiment | 2 | 0.82422 | 0.41211 | 12.1781 | 3.02E-05 | *** |
| Nematode inoculation | 1 | 0.00551 | 0.00551 | 0.1628 | 6.88E-01 | |
| Plant genotype | 3 | 0.11698 | 0.03899 | 1.1523 | 0.3344 | |
| Experiment x block | 9 | 0.04953 | 0.0055 | 0.1626 | 0.997 | |
| Nematode inoculation x Plant genotype | 3 | 0.03662 | 0.01221 | 0.3607 | 0.7816 | |
| Residuals | 68 | 2.30115 | 0.03384 | | | |
| **Tukey HSD** | | | | | | |
| | | **Experiment** | **P.adj** | **Signif.** | | |
| | | Summer 2021 - Winter 2020 | 0.882987 | | | |
| | | Winter 2021 - Winter 2020 | 0.000977 | *** | | |
| | | Winter 2021 - Summer 2021 | 6.51E-05 | *** | | |

*Table S2. Summary of the transcript, protein abundance and resistance outcomes in present study.*

| | **HG 0 IL w.r.t uninoculated** | | | | | **HG 2.5.7 IL w.r.t uninoculated** | | | | | **HG 1.3.6.7 IL w.r.t uninoculated** | | | | |
|---|---|---|---|---|---|---|---|---|---|---|---|---|---|---|---|
| | **Wm82** | **IL3849N** | **IL3849N +LC** | **IL3849N +HC** | **IL3025N** | **Wm82** | **IL3849N** | **IL3849N +LC** | **IL3849N +HC** | **IL3025N** | **Wm82** | **IL3849N** | **IL3849N +LC** | **IL3849N +HC** | **IL3025N** |
| **α-SNAP$_{Rhg1}$ LC mRNA** | NA | ↔ | ↓ | ↑ | NA | NA | ↑ | ↓ | ↑ | NA | NA | ↔ | ↔ | ↑ | NA |
| **α -SNAP$_{Rhg1}$ LC Protein** | NA | ↑ | **↑** | ↑ | NA | NA | ↑ | **↓** | ↑ | NA | NA | ↑ | **↔** | ↑ | NA |
| **α -SNAP$_{Rhg1}$ HC mRNA** | NA | NA | NA | **↓** | ↓ | NA | NA | NA | ↑ | ↑ | NA | NA | NA | **↓** | ↔ |
| **α -SNAP$_{Rhg1}$ HC Protein** | NA | NA | NA | **↑** | **↑** | NA | NA | NA | **↑** | **↑** | NA | NA | NA | **↑** | **↑** |
| **SHMT$_{Rhg4}$ mRNA** | ↑ | ↑ | ↑ | ↑ | ↔ | ↑ | ↑ | ↑ | ↑ | ↓ | ↑ | ↔ | ↓ | ↓ | ↔ |
| **SHMT$_{Rhg4}$ Protein** | ↔ | ↑ | ↑ | **↑** | **↑** | ↔ | ↔ | ↔ | ↔ | ↔ | ↔ | ↔ | ↔ | **↑** | **↑** |
| **NSF$_{RAN07}$ mRNA** | NA | ↑ | ↑ | ↑ | ↑ | NA | ↑ | ↑ | ↑ | ↔ | NA | ↑ | ↑ | ↑ | ↑ |
| **NSF$_{Total}$ Protein** | ↔ | ↔ | ↔ | ↑ | ↑ | ↔ | ↔ | ↔ | **↑** | **↑** | ↔ | ↔ | ↔ | ↑ | ↑ |
| **Avg. Female Index** | **100.0** | **4.07** | **2.01** | **1.74** | **21.36** | **100.0** | **13.04** | **8.13** | **2.01** | **75.01** | **100.0** | **76.90** | **67.46** | **71.31** | **25.30** |

For transcript abundance, large bold arrows represent statistically significant changes while small arrows indicate possible trends.

*Table S3. Primers used to quantify transcript abundances.*

| qPCR Primer Name | **5' to 3' Primer Sequence** | **Reference** |
|---|---|---|
| α-SNAP Forward | GCAATTGATGAAGAAGATGTTG | This study |
| α-SNAPRhg1 HC Reverse | TCAAGTAATAGCCTCATGCTG | |
| α-SNAPRhg1 LC Reverse | AATAACCTCATACTCCTCAAG | |
| SHMTRhg4 Forward | CTGCCATGACTTCTAGGGG | |
| SHMTRhg4 Reverse | AGCTTTGAGATCTTCAATAGCC | |
| NSFRAN07 Forward | CAACACGCCCGCGAGCGAC | |
| NSFRAN07 Reverse | GACCGCTGCCTATGGTG | |
| GmAAT Forward | GAGTTGATACCAGCCAACAG | (Beyer et al., 2021) |
| GmAAT Reverse | GAGACCAACAAGCAGTGTAG | |
| GmNIMIN1 Forward | ATGTTGAACACGGCATTCTC | |
| GmNIMIN1 Reverse | GGTACGGTGTGACTTTCTTG | |
| GmNREG-2 Forward | GTGTCTCGCATGTTTCATCC | |
| GmNREG-2 Reverse | GTGAGTATAGCAGCCACATC | |
